# The Curated Cancer Cell Atlas: comprehensive characterisation of tumours at single-cell resolution

**DOI:** 10.1101/2024.10.11.617836

**Authors:** Michael Tyler, Avishai Gavish, Chaya Barbolin, Roi Tschernichovsky, Rouven Hoefflin, Michael Mints, Sidharth V. Puram, Itay Tirosh

## Abstract

Single-cell RNA-seq (scRNA-seq) has transformed the study of cancer biology. Recent years have seen a rapid expansion in the number of single-cell cancer studies, yet most of these studies profiled few tumours, such that individual datasets have limited statistical power. Combining the data and results across studies holds great promise but also involves various challenges. We recently began to address these challenges by curating a large collection of cancer scRNA-seq datasets, and leveraging it for systematic analyses of tumor heterogeneity. Here we significantly extend this repository to 124 datasets for over 40 cancer types, together comprising 2,822 samples, with improved data annotations, visualisations and exploration. Utilising this vast cohort, we systematically quantified context-dependent gene expression and proliferation patterns across cell types and cancer types. These data, annotations and analysis results are all freely available for exploration and download via the Curated Cancer Cell Atlas (3CA) website (https://www.weizmann.ac.il/sites/3CA/), a central source of data and analyses for the cancer research community that opens new avenues in cancer research.

## Introduction

A tumour is a complex ecosystem of different cell types, genetic clones and dynamic cellular states. This intra-tumour heterogeneity (ITH) is central to tumour development and poses a major barrier to cancer therapy, with resistant tumour subpopulations driving continued disease progression^1^. Single-cell RNA-seq (scRNA-seq) has recently emerged as a powerful tool to study ITH, paving the way to a more complete understanding of cancer progression and treatment effects. Early studies applying scRNA-seq to patient tumour samples identified, for example, a stem/progenitor state in oligodendroglioma^2^, a partial epithelial-mesenchymal transition (pEMT) state in head and neck cancer^3^, and an antigen-presenting population of cancer-associated fibroblasts in pancreatic cancer^4^. Such discoveries were possible only with the high resolution and whole-genome coverage of scRNA-seq.

The generation of tumour scRNA-seq data has accelerated dramatically, with hundreds of recent publications. Collectively, the global cancer research community generated a vast pool of high-resolution tumour transcriptomic profiles^5,6^ that has the potential to transform our understanding of cancer and to drive the development of new treatment strategies. These datasets could ultimately define a foundational resource, replacing widely used bulk cohorts, such as The Cancer Genome Atlas (TCGA), with single-cell compendia that will be used routinely in cancer studies. However, due to cost and various technical constraints, individual scRNA-seq studies were only able to profile relatively few tumour samples, typically between 5 and 20. Each dataset is therefore severely underpowered to identify robust and clinically significant expression patterns. At the same time, the ability to compare data between studies is hindered by batch effects and inconsistencies in methods, formats and annotations.

We address these problems by curating a large number of published scRNA-seq datasets for combined analysis. We previously published a repository of 71 such datasets, in a pan-cancer scRNA-seq study characterising recurrent programs of transcriptional ITH^7^. We have now considerably expanded this cohort, almost doubling its size to 124 datasets, 2,822 samples and over 5.5 million single cells, enabling an even deeper exploration of ITH. We use this extended compendium to systematically identify context-dependent gene expression profiles, characterising cell type markers and identifying genes that distinguish malignant cells in various contexts. Furthermore, we present a comprehensive quantification of cell cycle, revealing high variability in proliferation rates across cell types and cancer types, and uncovering biases in cell cycle phases that are associated with driver mutations, most notably *TP53*. These data, analyses, and tools to explore them together constitute the Curated Cancer Cell Atlas (3CA), an updated resource which is available to the entire cancer research community via a website^8^ (https://www.weizmann.ac.il/sites/3CA/), and which enables comprehensive characterisation of tumours at single-cell resolution.

## Main

### Curating a comprehensive scRNA-seq data resource

To build 3CA, we conducted a thorough literature search to identify scRNA-seq cancer studies representing a wide range of cancer types, prioritising those having a relatively high number of samples, as well as smaller datasets for understudied cancer types. While the majority of these datasets were generated from patient samples, they also include cancer cell lines, organoids, mouse models and patient-derived xenografts (PDXs). Following this comprehensive search, we downloaded these datasets from their respective repositories, standardised their format and verified their cell annotations (**Fig. 1A**, upper half). An initial version consisting of 71 studies was reported previously^7^, and the updated version presented here consists of 124 datasets for over 40 cancer types, together comprising 2,822 samples and 5,536,628 cells.

**Figure 1.**
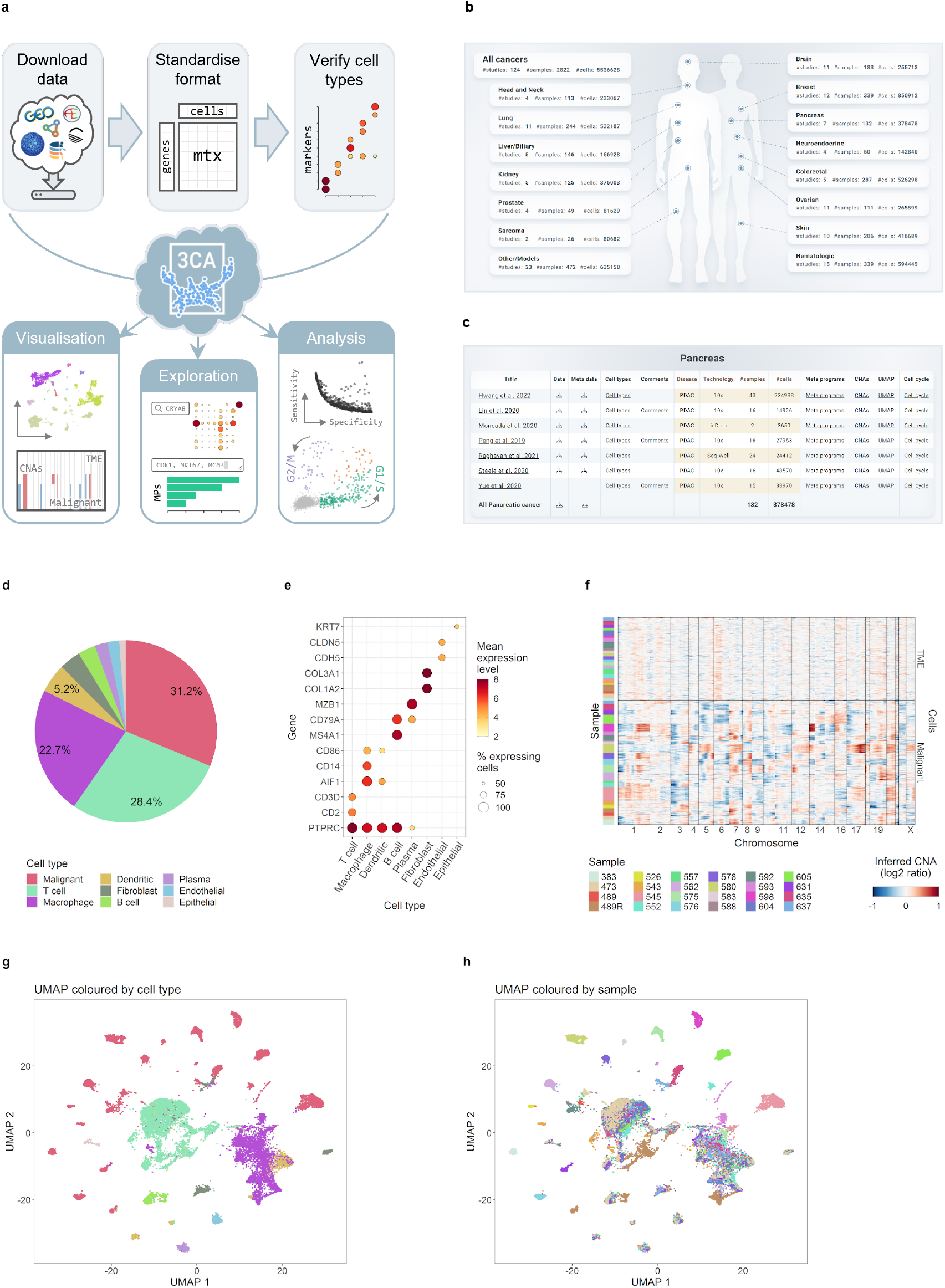
Overview of 3CA and data summary features. **a**. Scheme outlining the construction of 3CA and the features it contains. **b**. Screenshot of the 3CA website homepage, summarising the available datasets. **c**. Screenshot of the 3CA web page for pancreas, showing the available pancreatic cancer datasets with links and summary statistics. **d**. Pie chart showing the percentage of cells in the Raghavan et al. dataset^46^ assigned to each cell type. **e**. Dot plot showing the average expression level (colour) and percentage expressing cells (point size) of a selection of cell type marker genes (rows) in each cell type (columns) in the Raghavan et al. dataset^46^. **f**. Heatmap showing inferred copy number alteration (CNA) values (colour, quantified as log_2_ ratio, with blue indicating depletion and red amplification) at each chromosomal position (columns) for a representative subset of cells (rows) in the Raghavan et al. dataset^46^, with colour bar (left) showing the sample each cell belongs to. **g**. UMAP plot of all cells (points) in the Raghavan et al. dataset^46^, coloured by cell type, with the same colours as in **d. h**. UMAP plot as in **g** coloured by sample, with the same colours as in **f**.

Ensuring consistent annotations is especially crucial to data curation and enables combined analysis of this large cohort. We therefore standardised the cell type annotations and validated them in two ways. Firstly, we inferred copy number alterations (CNAs), using a method described previously^2,9^ (**Methods**), in order to confirm the annotation of malignant cells. Secondly, we verified the identity of the various non-malignant cell types in the tumour microenvironment by analysing the expression of canonical marker genes. Along with these cell annotations, each dataset’s sample metadata provides a variety of standardised sample annotations, including sample and patient IDs, disease name, sequencing technology and, when available, clinical annotations such as patient age, tumour stage and status of relevant driver mutations.

### An online portal for open access to 3CA

With its unparalleled size and rich annotations, 3CA is a valuable data resource for the entire cancer research community. To make it accessible to all researchers, we built and recently extended a website^8^ (https://www.weizmann.ac.il/sites/3CA/) via which the curated datasets are freely available to download, without the need for user registration or permissions. The home page summarises all datasets, organising them into 15 categories of cancer types. Separate category-specific pages contain links to download the expression matrices and/or sample metadata for the corresponding datasets, either per study or for all studies of this category at once (**Fig. 1B-C**).

We further sought to enrich 3CA with detailed data visualisations, functionality to explore the datasets, and new pan-cancer analyses (**Fig. 1A**, lower half). The category-specific pages of the website contain summary statistics for each dataset, including disease name, sequencing technology and cell and sample number, along with various visualisations for each dataset, including cell type composition, expression of canonical cell type marker genes, CNA matrices and UMAP plots coloured by cell type and sample (**Fig. 1D-H**). Additional website features are described below. Via this online portal, 3CA serves as a central source of data and analyses for all cancer researchers, which will continue to expand as the tumour scRNA-seq literature grows further.

### Exploring transcriptional tumor heterogeneity with 3CA

Beyond visualisation of the cell and sample composition of 3CA datasets, we added further features to the website that enable exploration of ITH at a higher level. We previously used 3CA for a pan-cancer characterisation of intra-tumor heterogeneity^7^. In particular, we identified recurrent programs of transcriptional ITH, which we term “meta-programs”. We defined a total of 149 meta-programs across 8 cell types, which collectively explain the majority of expression ITH. Thus, a given tumour sample may be described by quantifying the extent to which these meta-programs are variable across the cells in the tumour. We have therefore added to 3CA a summary, for each dataset, of the expression of meta-programs across cells of each type in each sample (**Fig. 2A-B**), as well as a feature for users to enter their own gene-sets and view their overlap with these meta-programs (**Fig. 2C**).

**Figure 2.**
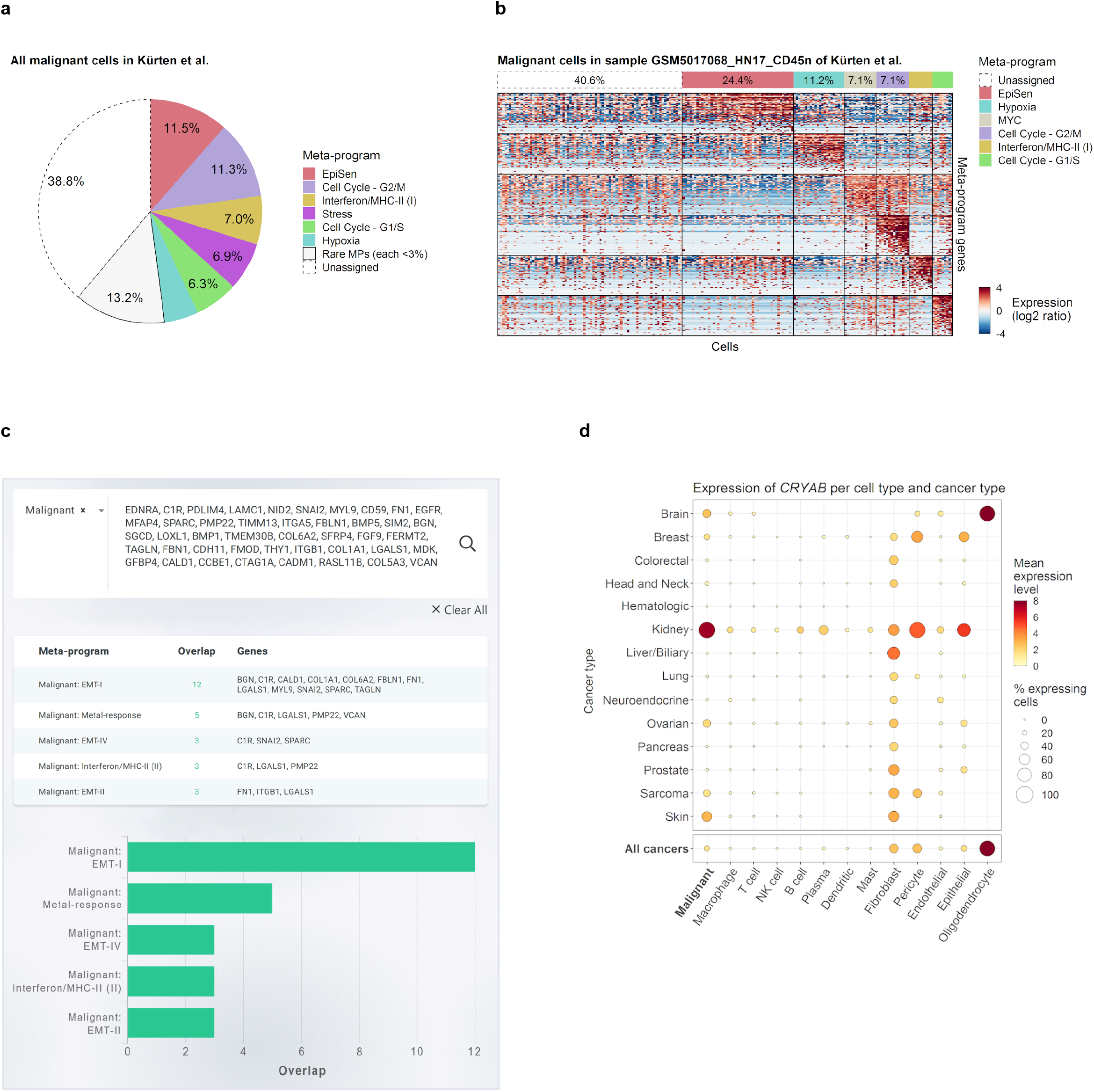
Data query features on the 3CA website. **a**. Pie chart showing the meta-program composition of malignant cells in the Kürten et al. dataset^47^. Each cell was assigned to at most one meta-program, with ambiguous cells classified as “Unassigned”. Percentages denote the proportion of cells in each category. **b**. Heatmap showing relative expression levels (colour, quantified as log_2_ ratio) of meta-program genes (rows) in malignant cells (columns) in sample GSM5017068_HN17_CD45n of the Kürten et al. dataset^47^. Each cell was assigned to at most one meta-program, with ambiguous cells classified as “Unassigned”. Colour bar (top) shows the classification of each cell, and percentages denote the proportion of cells in each category. **c**. Example output from the meta-program gene set query tool. Table shows, for each malignant cell meta-program having at least 2 genes in common with the input gene set, the size of its overlap with the input gene set and the genes residing in this overlap. Bar plot shows the size of the overlap (x axis) of each meta-program from the table (y axis) with the input gene set. **d**. Dot plot showing the average expression level (colour) and percentage expressing cells (point size) of *CRYAB* in each cell type (columns) and cancer type (rows, upper panel), and averaged across cancer types (bottom row).

We also added functionality to address one of the most common questions in cancer research, namely, given a gene of interest, what is its typical expression in each cell type and cancer type? Due to the resolution and comprehensiveness of 3CA, it offers the possibility to query the expression of any gene across many contexts. The 3CA website contains a search tool which returns, for any gene, a visualisation of its average expression, and the proportion of cells expressing it, per cell type and cancer type (**Fig. 2D**). This is further broken down per dataset to enable examination of study-specific effects.

### Characterising context-dependent gene expression patterns

The ability to resolve gene expression per cell type and cancer type also enables an unbiased search across genes to identify cases of highly context-dependent gene expression. This includes the characterisation of cell type markers, which we undertook for each of the most common cell types. This analysis, and those that follow, incorporated additional unpublished datasets on glioma, neuroendocrine tumours and head and neck cancer.

First, for a given cell type, we identified those genes whose average expression (across cancer types) was highest in this cell type. For each of these genes, we then defined a measure of its specificity and sensitivity as a marker of this cell type (**Methods**). Specificity reflects the extent to which this gene’s expression is unique to this cell type, while sensitivity reflects the likelihood of detecting expression of this gene in a cell of this type. These measures therefore assess how effectively an individual gene can identify the type of an individual cell. While this analysis is partially biased by the prior annotation of cell types in 3CA datasets, these annotations were generally determined at the level of clusters of cells and thus do not fully reflect marker performance at the level of individual cells.

Marker genes can be identified as those with unusually high specificity and/or sensitivity (**Fig. 3A**). An ideal marker should have both high specificity and high sensitivity, but due to the tradeoff between these measures, very few genes satisfy both of these criteria and it is important to assess both measures for each potential marker. Applying this approach to all cell types illuminated the distribution of markers across contexts (**Fig. 3B-C**).

**Figure 3.**
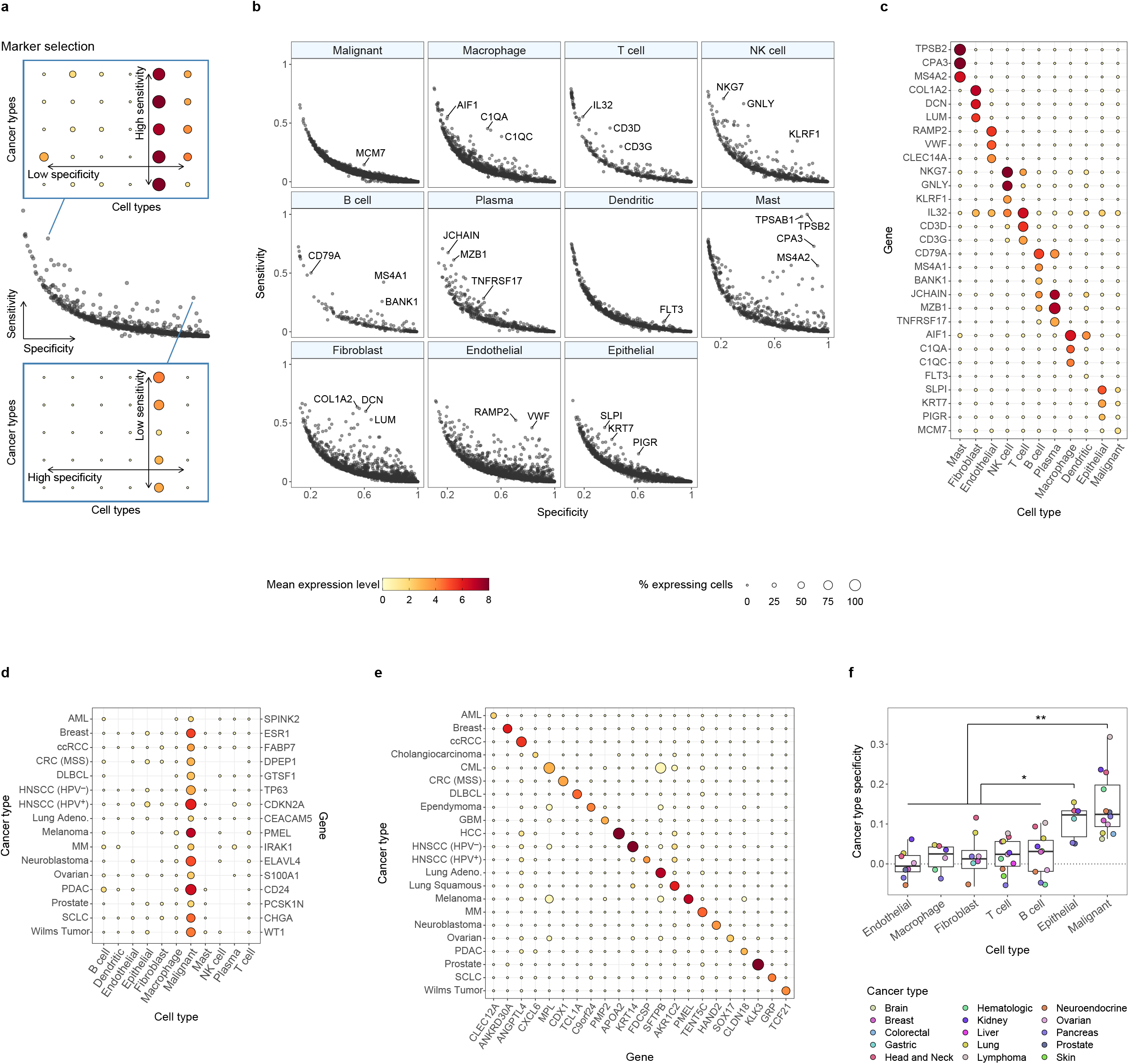
Context-dependency of gene expression. **a**. Scheme illustrating the selection of cell type marker genes. Middle panel (scatter plot) illustrates sensitivity (y axis) and specificity (x axis) of genes (points) for a given cell type, while the upper and lower panels (dot plots) illustrate expression levels (colour) and percentage expressing cells (point size) of example marker genes across cell types (columns) and cancer types (rows). **b**. Scatter plots per cell type showing sensitivity (y axis) and specificity (x axis) of genes (points) for each cell type. Selected genes with unusually high sensitivity or specificity are labelled. **c**. Dot plot showing the average expression level (colour) and percentage expressing cells (point size) in each cell type (columns) of marker genes labelled in **b** (rows). **d**. Dot plot showing the average expression level (colour) and percentage expressing cells (point size) in each cell type (columns) of selected malignant cell markers in each cancer type (rows). **e**. Dot plot showing the average expression level (colour) and percentage expressing cells (point size) of selected cancer-type-specific genes (x axis) in malignant cells in each cancer type (rows). **f**. Box plot showing the cancer type specificity (y axis) of each cell type (x axis) in each cancer type (points/colour). P values for the differences between cell types were computed by pairwise paired t tests, without adjustment. Asterisks indicate the maximal p value among pairwise comparisons between malignant cells, respectively epithelial cells, and all non-epithelial TME cell types, with ‘**’ and ‘*’ indicating max(p) < 0.01 and max(p) < 0.05 respectively. Pairwise differences between non-epithelial TME cell types, and between malignant and epithelial cells, are not significant.

Markers are strongest for mast cells, with multiple genes scoring exceptionally highly for both specificity and sensitivity (*TPSB2/AB1, CPA3, MS4A2*). In other cell types, however, the choice of markers represents a compromise between these two measures. For example, *NKG7* is a highly sensitive marker for NK cells, but it is not highly specific due to its expression in T cells. Conversely, *KLRF1* is highly specific to NK cells, but it is detected in only 60% of NK cells. Meanwhile, markers were unreliable for non-malignant epithelial and plasma cells, and all but absent for malignant and dendritic cells. The paucity of markers which are simultaneously sensitive and specific may be due to high context-specificity or high heterogeneity within a cell type. Malignant and epithelial cells are expected to have highly context-specific gene expression such that there is no universal epithelial or malignant marker. Dendritic cells divide into conventional and plasmacytoid, thereby precluding the existence of universal dendritic cell markers. Moreover, due to high expression similarities between certain cell types, such as between T cells and NK cells, some classical cell type markers have low specificity in scRNA-seq data (**Fig. 3B-C**).

As no pan-cancer markers were found for malignant cells, we instead focussed on identifying cancer-type-specific malignant cell markers. We found substantial heterogeneity in the strength and abundance of malignant markers between cancer types (**Fig. 3D, S1A**). Some markers were consistent with prior knowledge, including *PMEL* in melanoma, *CDKN2A* (encoding p16) in HPV^+^ head and neck cancer and *ESR1* in breast cancer (reflecting high representation of ER^+^ breast tumours in our cohort).

As well as malignant cell markers, we further characterised those genes whose expression in malignant cells is most variable between cancer types (regardless of their cell type specificity). This analysis identified many highly context-dependent genes, with one of the most prominent being *KLK3*, encoding prostate-specific antigen (PSA), which is highly specific to prostate cancer (**Fig. 3E, S1B**). Other examples of cancer-type-specific genes included *APOA2* in liver cancer and *ANKRD30A* in breast cancer.

### Cancer type specificity is unique to malignant and epithelial cells

While we could find many genes with robust cancer-type-specific expression among malignant cells (**Fig. 3E**), we could find far fewer for TME cell types (**Fig. S2**). To better characterise the context specificity of gene expression in different cell types, we aggregated the cells of each cell type from each tumor into a pseudobulk profile, and then compared these pseudobulk profiles across all tumors. This analysis revealed extensive diversity for each cell type, reflecting the combined effects of cancer type, cancer subtype and genetics, the TME and spatial location, technical batch effects and possibly other effects. This complex set of effects may be explored further in future work, while here we focus on only quantifying the effect of cancer type while controlling for technical batch effects. We defined an overall expression similarity between every pair of pseudobulk samples and then classified the pairs based on whether they are from the same cancer type and whether they are from the same study (i.e. batch). The effect of cancer types can be quantified by the average similarity of pairs from the same cancer type vs. the average similarity of pairs from different cancer types. To control for batch effects, both measures of similarity were calculated only for pairs from different studies (**Fig. S3A**).

Surprisingly, for most cell types we observed comparable similarity of pairs of pseudobulk samples from the same or from different cancer types, indicating a minimal effect of cancer type (**Fig. 3F**). The only exceptions were the malignant cells, followed by non-malignant epithelial cells. Malignant cells from different cancer types are associated with distinct sets of common genetic events and thus are expected to be highly distinct. The non-malignant epithelial cells reflect parenchymal cells that are also expected to vary considerably between tissues. Yet all the other TME immune and stromal cell types appear to have very limited cancer type specificity. A similar, though weaker, effect was observed for patient specificity of cell types, though this analysis does not control for batch effects (**Fig. S3A-B**). Thus, immune and stromal cell types are diverse even within an individual tumor and are affected by various processes, but their overall expression profile within tumors appears to be only minimally dependent on the cancer type. This observation supports the pan-cancer approach for discovery of cellular states in immune and stromal cell types^10–13^.

### Pan-cancer comparison of proliferation rates

Proliferation is a defining feature of cancer, but the cell cycle behaviour of different tumour types remains poorly characterised. Dozens of canonical cell cycle genes are highly upregulated during the cell cycle, in a phase-dependent manner^2,14^. Hence, scRNA-seq allows an efficient means to detect cycling cells and assign them to specific phases. Accordingly, 3CA offers an unprecedented opportunity to systematically compare cell cycle patterns across contexts.

We quantified cell cycle across datasets by scoring cells for G1/S and G2/M gene signatures and defining appropriate thresholds (**Fig. 4A, Methods**). Summary plots of these cell cycle measurements are available for each published dataset on the 3CA website. The gene signatures were adapted for each dataset, but highly similar results were obtained when using the same consensus signatures for all datasets (**Fig. S4**). Sequencing technology also had negligible effect on downstream results (**Fig. S5**).

**Figure 4.**
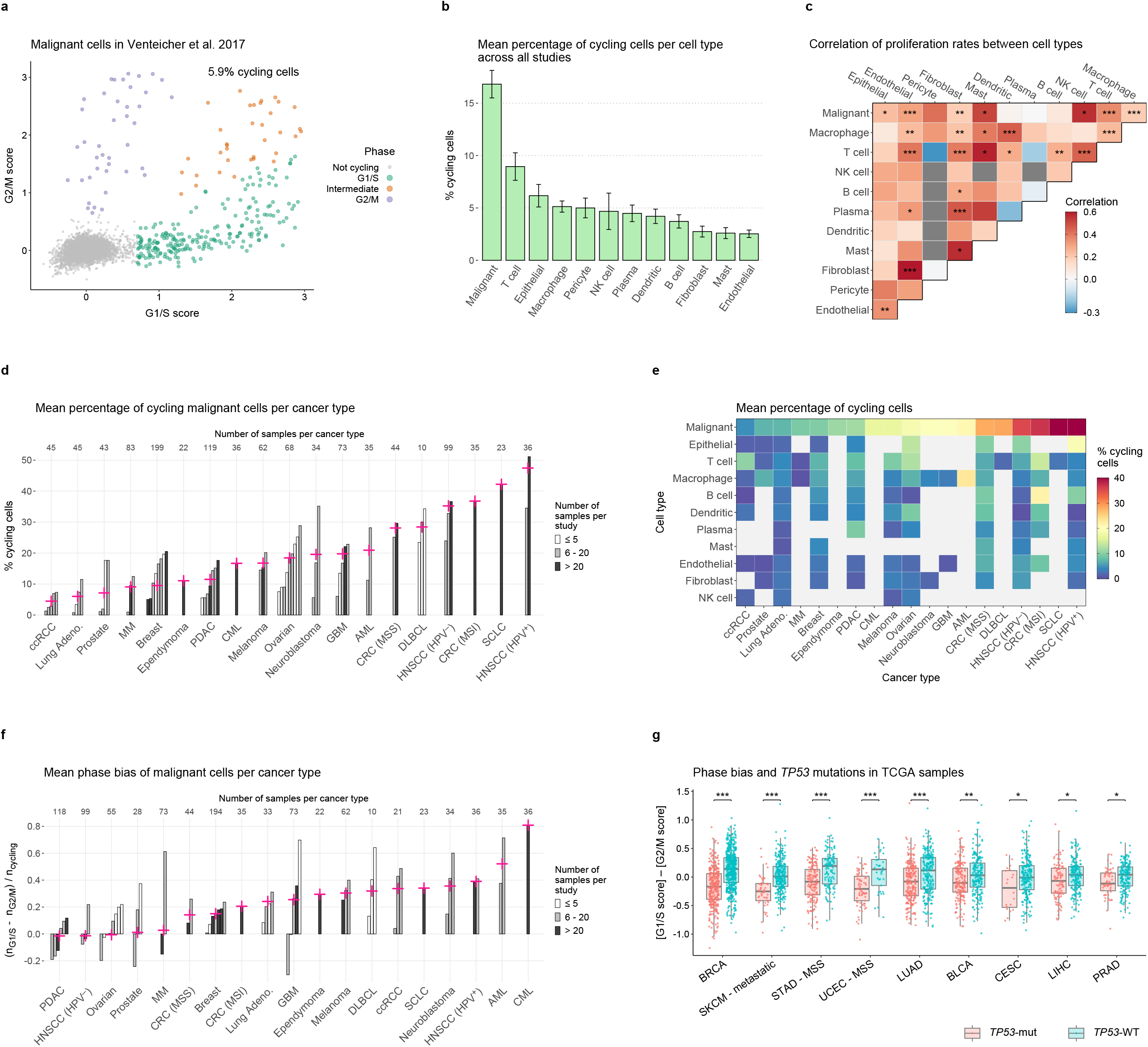
Quantification and comparison of cell cycle patterns. **a**. Scatter plot showing scores for gene signatures of G1/S (x axis) and G2/M (y axis) for all malignant cells (points) in the Venteicher et al. dataset^48^, coloured by cell cycle phase. **b**. Bar plot showing the average percentage of cycling cells (y axis, average across all studies) in each cell type (x axis). Error bars denote the standard error. **c**. Heatmap showing the Spearman correlation between cell types (colour, correlation computed across samples) of percentages of cycling cells. Asterisks indicate the statistical significance of the correlation (computed via algorithm AS 89^43^), after adjusting to FDR < 0.05, with ‘***’, ‘**’ and ‘*’ indicating p < 0.001, p < 0.01 and p < 0.05 respectively, and no asterisk indicating not significant. **d**. Bar plot showing the percentage of cycling malignant cells (y axis) in each study (bars), grouped by cancer type (x axis), with crosses denoting the average y value for each cancer type, weighted by the number of samples in each study which contain at least 10 malignant cells. Bar colour categorises studies by number of such samples, and values above the plot denote the total number of such samples per cancer type. **e**. Heatmap showing the weighted average of the percentage of cycling cells (colour, defined as for the crosses in **d**) per cancer type (x axis) and cell type (y axis). Grey squares indicate insufficient data (missing values). Cell types are ordered by the average of their non-missing values, and cancer types are ordered by the values for malignant cells. **f**. Bar plot showing the phase bias (y axis, quantified by the relative fraction of cycling cells in G1/S versus G2/M) of malignant cells in each study (bars), grouped by cancer type (x axis). Crosses, bar colour and number of samples per cancer type are defined as in **d**. Low and high y values indicate relative bias toward G2/M and G1/S, respectively. **g**. Box plot showing the phase bias (y axis, quantified by difference between scores for G1/S and G2/M gene signatures) of TCGA tumour samples (points), coloured by *TP53* mutation status, for cancer types having a significant difference in y values between *TP53*-mut and *TP53*-WT tumours. Significance values were computed by t test and adjusted to FDR < 0.05. Asterisks indicate significance after adjustment in each case, with ‘***’, ‘**’ and ‘*’ indicating p < 0.001, p < 0.01 and p < 0.05 respectively. Low and high y values indicate bias toward G2/M and G1/S, respectively.

A comparison of proliferation rates between cell types (**Fig. 4B**) confirmed malignant cells as the most proliferative, with more than 15% of malignant cells typically observed cycling.

However, all cell types show some cycling activity, with surprisingly high proliferation in non-malignant cells, especially T cells and normal epithelial cells. Moreover, we observed an overall positive correlation of proliferation rates between cell types across tumors (**Fig. 4C, S6**), consistent with recent findings^7^. This suggests that cell cycle may be stimulated in multiple cell types at once by microenvironmental factors and intercellular communication. An especially high correlation was observed between proliferation of fibroblasts and endothelial cells.

Proliferation rates were highly variable between cancer types, especially for malignant cells (**Fig. 4D-E, S7**). Proliferation of malignant cells is lowest in clear cell kidney cancer (∼5% cycling cells), consistent with the slow growth of kidney tumours and their resistance to chemotherapy^15^. At the opposite extreme, HPV^+^ head and neck cancer is the most highly proliferative cancer type (>45% cycling cells). This may be explained by the mechanism of action of HPV, which silences p53 and pRb activity to promote progression through the cell cycle^16^. However, HPV^−^ head and neck cancer is also among the most highly proliferative cancer types (∼35% cycling cells), suggesting that high proliferation is a general feature of head and neck cancer.

### Variability in phase bias explained by specific genomic alterations

As our method for measuring cell cycle state distinguishes G1/S and G2/M phases, we were able to explore patterns of phase bias, that is, the relative proportion of cycling cells detected in G1/S versus G2/M. Technical confounders, such as the choice of gene signatures and sequencing technology, had negligible effect on our estimates of phase bias and downstream results (**Figure S4** (right panel), **S5D**). As with overall proliferation, we observed high variability in phase bias across cancer types, especially in malignant cells, with acute and chronic myeloid leukaemias being the most strongly biased toward G1/S, and pancreatic and HPV^−^ head and neck cancers having the strongest relative bias toward G2/M (**Fig. 4F, S8**). Interestingly, while HPV^+^ and HPV^−^ head and neck cancers are both among the most proliferative cancer types overall (**Fig. 4D-E**), they had opposite patterns of phase bias, with HPV^+^ exhibiting strong bias toward G1/S.

To test whether genomic alterations may explain the variation in phase bias across cancer types, we also computed phase bias scores in bulk RNA-seq profiles from TCGA for a variety of cancer types (**Methods**). Using these scores, we observed a strong association between G2/M bias and *TP53* mutations in multiple cancer types, consistent with the role of p53 as a gatekeeper of the G1/S transition (**Fig. 4G, S9A**). Moreover, an unbiased analysis of many genes commonly mutated in cancer^17^ identified *TP53* mutations as the most consistently associated with G2/M bias across cancer types (**Fig. S9B**). Interestingly, this analysis suggested *RB1* mutations as the most consistently associated with G1/S bias (**Fig. S9B-C**). As HPV acts in part through degradation of pRb^18^, which is not mutated in HPV^−^ head and neck tumours, this association could explain the opposite phase bias patterns observed in HPV^+^ and HPV^−^ head and neck tumours. Various other driver mutations have strong context-specific associations with phase bias, such as *SMARCA4* and *EGFR* in lung adenocarcinoma, *PIK3CA* and *CDH1* in breast and stomach cancers, and *CTNNB1* in endometrial cancer. Together, this analysis indicates that phase bias in cycling malignant cells is influenced by various cancer driver alterations, including those affecting *TP53* and *RB1*.

## Discussion

3CA brings together many individual scRNA-seq efforts from across the cancer research community to unlock their combined potential. The volume and variety of curated data, along with its website availability, confers a level of accessibility and statistical power to scRNA-seq that was previously lacking in cancer research. This data resource will be immensely valuable to many research groups for various tasks, such as: (i) to search for and download individual datasets most suited to a particular question; (ii) to examine expression patterns of genes of interest across cell types and cancer types; (iii) to examine the expression of known gene signatures in new contexts and to test their robustness and generality; (iv) to conduct various pan-cancer analyses and uncover relationships between diseases; and (v) to fine-tune statistical models and algorithms. 3CA fills the role of a central source of scRNA-seq data for all cancer researchers, who may in turn contribute new datasets to further enrich this resource.

While other studies have presented repositories for processed scRNA-seq data^19–24^, these either have limited size, lack a cancer focus, or are centred around tumour microenvironmental factors such as immune cells. 3CA prioritises malignant cells and contains extensive data visualisations and analyses detailing their diversity within and between cancer types. Our efforts also included careful curation of tumour clinical annotations, which are typically sparse in individual scRNA-seq datasets. As 3CA continues to grow, these clinical annotations will enrich future research efforts by uncovering single-cell expression patterns that correlate with clinical outcomes.

In our processing and analysis of 3CA datasets, we avoided using data integration methods, such as those offered by scANVI^25^, Harmony^26^ and Seurat^27^. These methods attempt to remove batch effects in scRNA-seq data using certain assumptions as to what constitutes technical artefacts, but there is currently no widely agreed standard, and they will likely remove some true biological signal^28^. This is especially true in the context of cancer, where much of the transcriptional variation between tumour samples arises from their unique genetic and epigenetic profiles, rather than from batch effects. Importantly, our analysis has either focused on intra-tumour heterogeneity, with comparisons between samples at the level of gene signatures rather than expression levels, or has reported only average values across many samples. The high sample size in 3CA allows confidence in these average values, and their accuracy will improve further with the inclusion of more datasets. However, there would be clear advantages to a fully integrated scRNA-seq data resource of this size, in which expression levels may be directly compared between any two samples. Further research is needed to establish optimal methods to integrate 3CA data while preserving biological signal.

We anticipate the further extension of 3CA in at least three ways. Firstly, sequencing efforts are becoming larger, with individual studies sequencing upwards of 50 samples. Including more such studies will significantly boost 3CA’s statistical power. Secondly, we expect new studies to include cancer types currently underrepresented in 3CA, as well as rare sample types such as post-treatment tumours, metastatic lesions and circulating tumour cells (CTCs). Thirdly, single-nucleus RNA-seq from frozen tumour samples is quickly becoming common^29–34^, and methods also exist for profiling fixed tissue^35–39^. These technologies open up many new possibilities by lifting the restriction to sequencing fresh tissue, and hence we expect 3CA to grow and diversify substantially in the coming years.

## Methods

### Data preprocessing

For all analyses in this study, datasets were filtered to remove cells with fewer than 1000 detected genes. Following this, expression levels were converted to log_2_(TPM/10 + 1), where TPM denotes transcripts per million, and the factor 1/10 reflects an estimated upper bound of 100,000 for the number of transcripts in single cell libraries.

### Data visualisations on the 3CA website

To generate UMAP plots for each dataset, we removed samples having fewer than 10 cells, then restricted to genes for which log_2_(mean(TPM) + 1) ≥ 4. A partial singular value decomposition was computed using the IRLBA algorithm^40^, and a UMAP was computed using the first 50 of the resulting principal components. Visualisations of cell type marker expression in each dataset were made using a manually curated list of canonical cell type markers, and restricting to cell types having at least 10 cells in the given dataset.

Copy number alterations (CNAs) were inferred from the scRNA-seq data using a method which was described previously^2,9^, and which follows the approach of the infercna R package (https://github.com/jlaffy/infercna). Briefly, in each dataset, we ordered the genes by chromosomal position, then computed the running average of their centred expression levels. After median-centring per cell, we adjusted these values relative to those of a selection of reference cells. These reference cells were chosen to be confidently nonmalignant, and to have expression profiles as similar as possible to those of the presumed malignant cells. The exact choice of reference cells was specific to each dataset, depending on the cell types detected therein.

### Meta-program exploration features of the 3CA website

To measure the distribution of meta-program expression among cells of a given type in each sample or dataset, we computed meta-program scores using a method described previously^2^, which measures the expression of signature genes relative to a set of control genes. These control genes are chosen to have comparable expression levels to the signature genes but no coherent association with any biological process. Briefly, genes were ranked by average expression and divided into a number of discrete bins. Then, for each signature gene *g*, we sampled a set of control genes from the corresponding expression bin, and computed the difference between the expression of *g* and the average expression of these control genes. The signature score was then defined as the average of these relative expression levels across all genes in the signature.

Having scored the cells for each of the meta-programs for the corresponding cell type, we assigned each cell to a meta-program as follows. If none of a cell’s meta-program scores was greater than 1, this cell was classified as “unassigned”, while if at least one score was greater than 1, the cell was assigned to the meta-program with maximal score. Among the cells given an assignment, rare meta-programs (with “rarity” depending on whether this analysis was per-sample or per-dataset) were considered spurious, and cells assigned to them were reclassified as “unassigned”.

### Average gene expression per cell type, study and cancer type

For a given gene, and for each of expression level and percentage expressing cells, the average for a given cell type and study or cancer type was computed as follows. First, we computed the average across all cells of that type from a given study that were profiled with the same sequencing technology. If a study used more than one sequencing technology, these values were further averaged across technologies within the study. We then calculated the weighted average of these values across datasets for each cancer type, with weights equal to the number of samples in each dataset having at least 10 cells of that type.

### Sensitivity and specificity of gene expression

We included in this analysis only those cancer types having at least 2 studies and 10 samples in total, or one study with at least 20 samples. We included genes that were detected in datasets for at least 20 cancer types. Then, for each gene, we conducted three comparisons of its average expression levels (average per cell type and cancer type, defined above) to define sensitivity and specificity to three different contexts:

i. For the first comparison, we first computed the median of the given gene’s averages for each cell type (median across cancer types), then compared these medians between cell types. The values of interest are therefore the cell type medians, and the contexts for comparison are the different cell types, with the context of interest being a single chosen cell type.
ii. For the second comparison, we first fixed a cancer type, then compared the given gene’s averages between malignant cells and all other cell types within this cancer type. The values of interest are the gene averages, and the contexts for comparison are the cell types within this cancer type, with the context of interest being the malignant cells. This comparison was repeated with each cancer type, excluding carcinomas with insufficient data for non-malignant epithelial cells.
iii. For the third comparison, we first fixed a cell type (namely malignant cells), then compared the given gene’s averages between malignant cells from different cancer types. The values of interest are the gene averages, and the contexts for comparison are malignant cells in different cancer types, with the context of interest being malignant cells in a single chosen cancer type.

In each case, if the relevant value was highest in the context of interest, its sensitivity to this context was defined as this value (the highest value across contexts), and its specificity was defined as 1/(*x* + 1), where *x* is the second-highest value across contexts. The sensitivity measures were then scaled to the interval [0, 1] by dividing by the maximum across genes.

### Number of cancer-type-specific genes per cell type

For each cell type, we quantified the number of cancer-type-specific genes for this cell type as follows. For each gene whose average expression was highest in this cell type, we defined a value *y* to be this average value (corresponding to the sensitivity described above), and a value *x* to be its highest average expression level among all other cell types (corresponding to the value *x* described above). Then, for each value *c* from a manually selected set of thresholds, we defined the number of cancer-type-specific genes for this cell type to be the number of genes with *y* > *x* + *c*. As expression levels were defined in log space, *c* = 1, respectively *c* = 2, 3 identifies genes with ≥2-fold, resp. 4-, 8-fold higher expression in this cell type than in other cell types.

### Cancer type and patient specificity of cell types

In datasets having at least 11,000 measured genes, pseudo-bulk profiles were computed for each cell type in each tumour sample by averaging the expression levels (transcripts per 100,000) for each gene across cells of this cell type in this sample. We further excluded genes measured in fewer than 80% of these datasets, then, after excluding ribosomal genes, restricted to the top 6000 genes with highest average expression across pseudo-bulk profiles. We then log-transformed these profiles and computed their pairwise Pearson correlations. Cancer type specificity was then defined, for each cell type and cancer type, by the average correlation between pseudo-bulks of this cell type and cancer type from different studies, minus the average correlation between pseudo-bulks of this cell type from different cancer types. Patient specificity was defined as 1 minus the average correlation between pseudo-bulks of this cell type from the same study. Differences between cell types were assessed using pairwise paired t tests. P values were not adjusted.

### Cell cycle quantification and phase assignment

To analyse the expression of cell cycle genes in each dataset, we constructed initial G1/S and G2/M gene signatures by taking the unions of published G1/S, respectively G2/M signatures from the Scandal R package (https://github.com/dravishays/scandal) and 4 other sources^2,7,41,42^. We then removed genes in the overlap of the resulting G1/S and G2/M gene sets. G1/S and G2/M scores were defined similarly to the meta-program scores described above, with slight differences. For a given signature gene *g* and corresponding set of control genes, we assigned two values to *g*. First, we computed the difference between the expression level of *g* and the average expression level of the sampled control genes, capping these differences at +/-3 to lessen the influence of extreme values. Second, we assigned 1 if the expression level of *g* was greater than the bin average, and 0 otherwise. We then defined two scores for each gene signature by averaging each of these sets of values across all genes in the signature. We refer to these two scores as “mean-based” and “count-based”, respectively.

In each dataset, we then filtered the G1/S and G2/M signatures to maximise orthogonality. To do this, we computed the correlation of each gene with the mean-based G1/S and G2/M scores, and excluded genes for which the difference between these correlations was 0.1 or less. If, after this step, there were fewer than 20 genes remaining in either signature, this dataset was excluded from further analysis. If there were more than 50 genes remaining in a given signature, we ranked them by their correlation values for the same signature, and by the differences between their G1/S and G2/M correlations. We retained the top 50 genes according to the average of these ranks. Note this procedure meant that we obtained different filtered signatures for each dataset. We further defined consensus signatures by ranking all G1/S and G2/M genes by their occurrence in the dataset-specific signatures, retaining the top 50. We compared the cell cycle estimates yielded by the consensus and dataset-specific signatures (**Fig. S4**), but all downstream scRNA-seq analysis used the dataset-specific signatures.

After recalculating the G1/S and G2/M scores using these filtered signatures, we defined thresholds to distinguish cycling from non-cycling cells using a bootstrapping approach. For each cell type, we generated 1000 null distributions consisting of mean- and count-based scores for 100 “pseudo-cells”. These scores were computed as above, except that for each signature gene, expression levels were taken from a different set of 100 cells that were randomly sampled (with replacement) from the expression matrix. This approach removed the correlation between signature genes while preserving each gene’s distribution of expression levels. Then, for each cell *c* and each score type, a p value was obtained by one-sided binomial test on the number of null distributions containing the score for *c*, i.e. in which the highest-scoring pseudo-cell scored higher than *c*. After adjustment to FDR < 0.05, cells were classified as cycling if, for both score types, the p value for either G1/S or G2/M score was less than 0.05. To correct for biases in the thresholds due to high proportion of cycling cells, for cell types of which more than 10% were classified as cycling, we reassigned each signature gene to a control gene set based on its average expression in the non-cycling cells. Scores, null distributions, p values and cycling assignments were then re-computed for these cell types as before.

For each of G1/S and G2/M signatures, we next defined a consensus significance variable, taking the value 1 if adjusted p values for both mean- and count-based scores were less than 0.05, and 0 otherwise. We then defined new mean-based score thresholds in each cell type by fitting a binomial regression model of the mean-based score against this consensus significance. Cells were re-classified as cycling if their mean-based scores for either G1/S or G2/M passed the corresponding regression threshold. Scores were then re-centred per cell type relative to the average mean-based scores of the non-cycling cells. Final mean-based score thresholds for each of G1/S and G2/M were defined by manually examining the distribution of the regression thresholds across all cell types and datasets and choosing an appropriately conservative consensus value. Cells were reclassified as cycling according to these consensus thresholds. Lastly, cells were assigned to G1/S and G2/M phases using a fold change threshold of 2 on re-centred mean-based scores. Cells assigned to neither phase were classified as intermediates.

### Estimates of cell cycle proportion and phase bias

Estimates of proportion of cycling cells and phase bias were computed per study, sequencing technology, cancer type and cell type only for those combinations having at least 100 cells. For each such combination, to avoid bias arising from samples with exceptionally many or few cycling cells, we defined outlier samples as those with number of cells outside the bounds [25^th^ percentile – 1.5×IQR, 75^th^ percentile + 1.5×IQR]. Samples below the lower bound were excluded from further analysis, while those above the upper bound were down-sampled to this bound. The proportion of cycling cells across these samples was then defined as the number of cells classified as cycling (of any phase) divided by the total number of cells. If there were at least 100 cycling cells across these samples, the phase bias was defined as the difference between the number of cells assigned to G1/S and the number assigned to G2/M, divided by the total number of cycling cells. If a study contained data generated by multiple sequencing technologies, the estimates for this study were averaged across sequencing technologies, weighted by the number of samples containing at least 10 cells, to obtain one estimate per study, cancer type and cell type for each of cycling proportion and phase bias. To obtain pan-cancer cell-type-level estimates, these study-level estimates were averaged across all studies, weighted by number of samples with ≥10 cells. To obtain cancer-type-level estimates for each cell type, the study-level estimates were averaged across studies, weighted by number of samples with ≥10 cells. Cycling proportions were also calculated per sample, only in those samples having at least 100 cells. These computations were also performed after restricting to 10x datasets.

### Correlation of cell cycle between cell types

To compute the correlation of cell cycle between cell types across all cancer types, we first centred the per-sample estimates of cycling proportion within each study and cell type. Then, for each pair of cell types having such estimates in at least 10 of the same samples, Spearman correlation was computed between cycling proportions across these common samples. We computed the correlation of cell cycle between cell types within each cancer type, and the significance of these correlations, using the same procedure per cancer type, before filtering out cell types having fewer than 20 such correlation values across all cancer types and cell type pairs. Significance levels were computed via algorithm AS 89^43^ and adjusted to FDR < 0.05.

### TCGA mutations analysis

TCGA expression and mutation data were obtained from the Broad GDAC Firehose (gdac.broadinstitute.org). The expression data was downloaded in the form of RSEM^44^ “scaled estimates”, then multiplied by 10^6^ (giving a measure similar to transcripts per million) and log-transformed. For colorectal and stomach cancers, MSI tumours were defined as those with more than 500 mutations in total, and similarly, MSI endometrial tumours were defined as those with more than 400 mutations. Following this, we restricted our attention to a previously defined list of genes commonly mutated in cancer^17^, and among these, we focussed on nonsense, nonstop, frame shift and splice or translation start site mutations, along with missense mutations and in-frame insertions/deletions occurring in at least 2 tumours.

Cell cycle scores were computed per cancer type using the consensus cell cycle gene signatures, via the method described above, with bulk profiles in place of individual cells. We then defined the phase bias as the difference between G1/S and G2/M scores. In each cancer type having at least 100 tumours, and for each gene from the above list which was mutated in at least 10 tumours and at least 5% of tumours, and wild type in at least 10 tumours, we compared phase bias scores between mutated and wild type tumours via two-sample t tests. P values were adjusted to FDR < 0.05 separately for each of *TP53* and *RB1*.

## Data Availability

This study used only external datasets and did not involve the generation of new data. All published single-cell datasets are available on the 3CA website^8^ (https://www.weizmann.ac.il/sites/3CA/), with the exception of one dataset^45^, for which permission for sharing through 3CA was not granted, and which is available through EGA with accession number EGAS00001002543. Additional unpublished datasets used will be added to the 3CA website when possible. TCGA data was downloaded from http://gdac.broadinstitute.org/.

## Code Availability

Source code for all analyses in this study, and for generating figures available on the 3CA website, will be made available on Github at https://github.com/tiroshlab/3ca.

## Acknowledgements

We thank all members of the Tirosh lab for helpful discussions and comments. This work was supported by the Israeli Council for Higher Education (CHE) via the Weizmann Data Science Research Center, and by the Israel Science Foundation. I.T. is the incumbent of the Dr. Celia Zwillenberg-Fridman and Dr. Lutz Zwillenberg Career Development Chair, and is supported by the Zuckerman STEM Leadership Program, the Mexican Friends New Generation, and the Benoziyo Endowment Fund.

