## Supplementary Figures for "The Curated Cancer Cell Atlas: comprehensive characterisation of tumours at single-cell resolution"

**a**

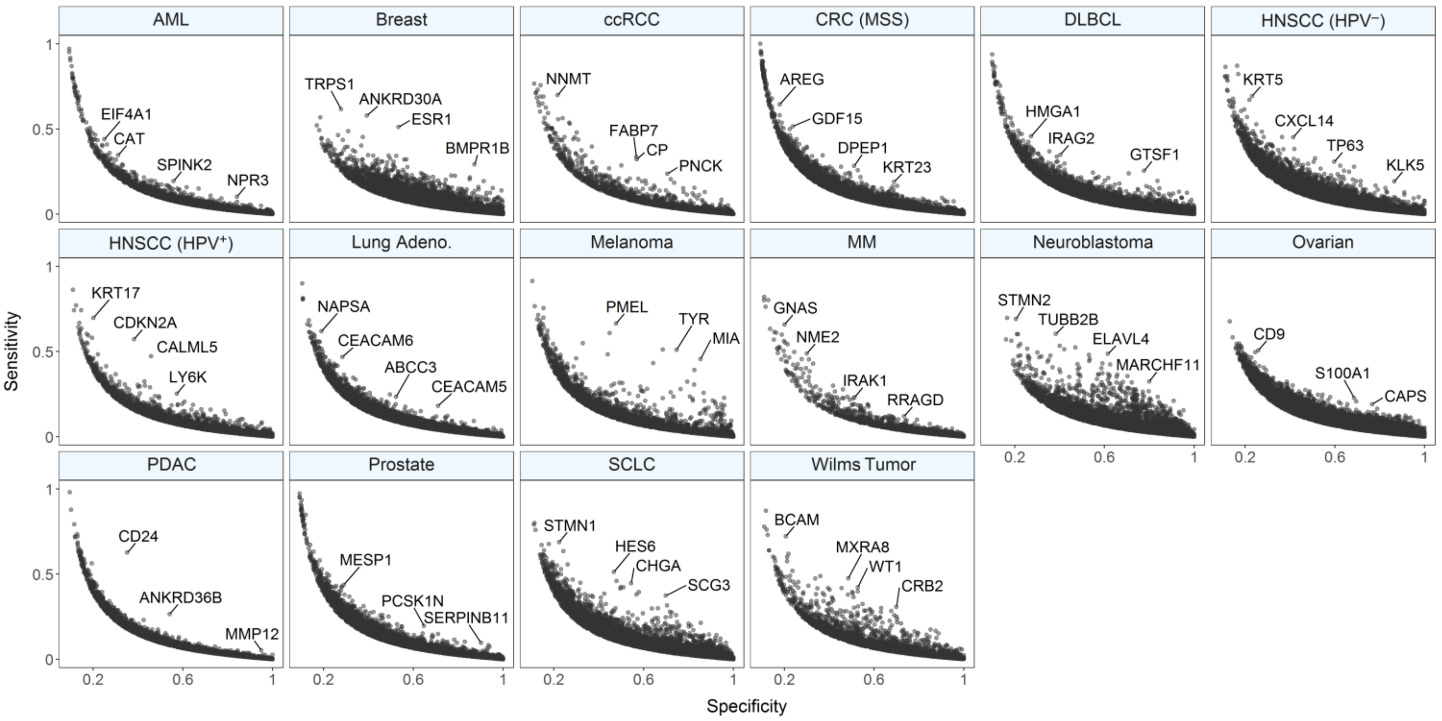

AML
Breast
ccRCC
CRC (MSS)
DLBCL
HNSCC (HPV⁻)
HNSCC (HPV⁺)
Lung Adeno.
Melanoma
MM
Neuroblastoma
Ovarian
PDAC
Prostate
SCLC
Wilms Tumor
EIF4A1
CAT
SPINK2
NPR3
TRPS1
ANKRD30A
ESR1
BMPR1B
NNMT
FABP7
CP
PNCK
AREG
GDF15
DPEP1
KRT23
HMGA1
IRAG2
GTSF1
KRT5
CXCL14
TP63
KLK5
KRT17
CDKN2A
CALML5
LY6K
NAPSA
CEACAM6
ABCC3
CEACAM5
PMEL
TYR
MIA
GNAS
NME2
IRAK1
RRAGD
STMN2
TUBB2B
ELAVL4
MARCHF11
CD9
S100A1
CAPS
CD24
ANKRD36B
MMP12
MESP1
PCSK1N
SERPINB11
STMN1
HES6
CHGA
SCG3
BCAM
MXRA8
WT1
CRB2
Sensitivity
Specificity
0
0.5
1
0.2
0.6
1


**b**

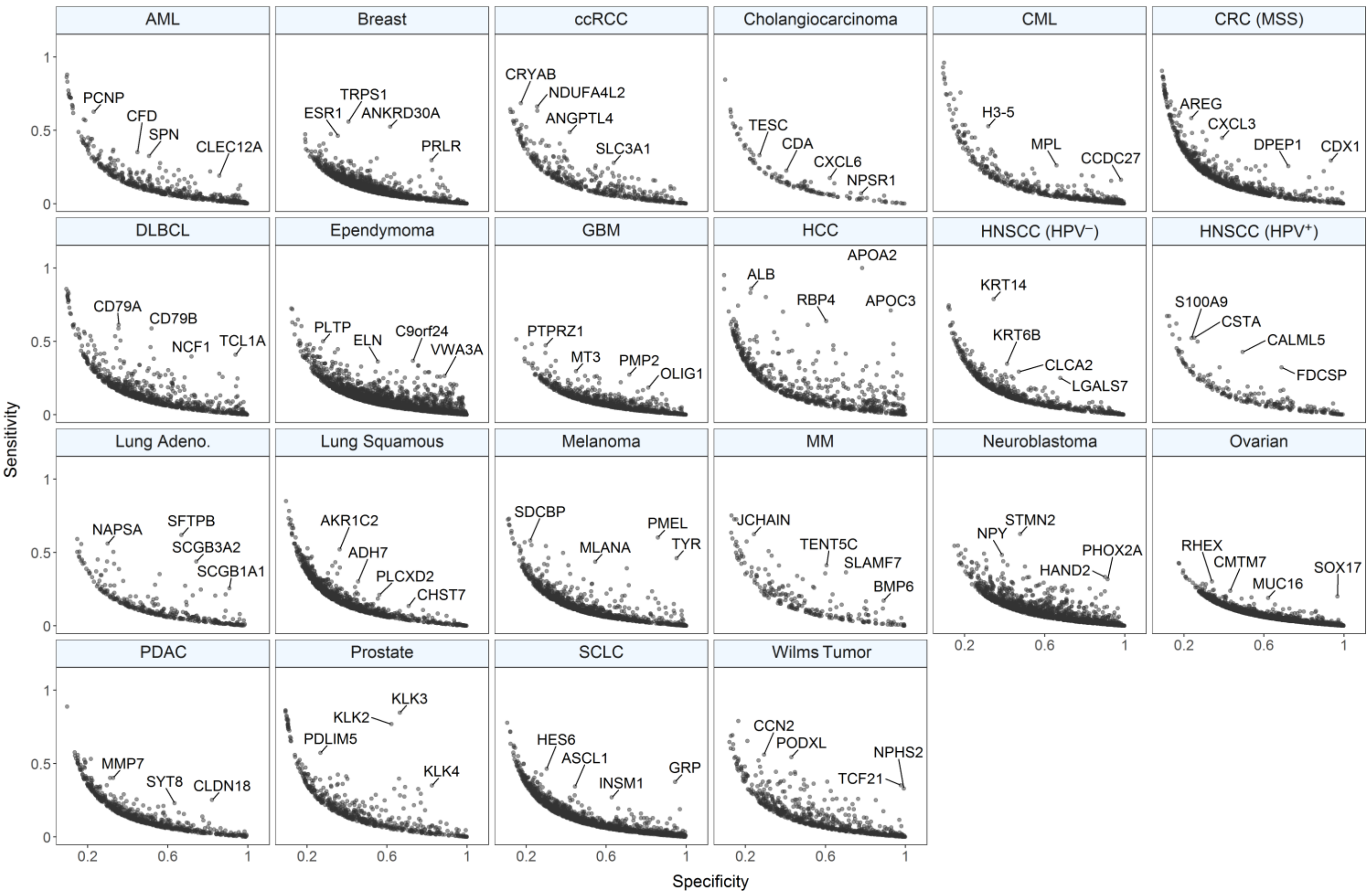

AML
Breast
ccRCC
Cholangiocarcinoma
CML
CRC (MSS)
DLBCL
Ependymoma
GBM
HCC
HNSCC (HPV⁻)
HNSCC (HPV⁺)
Lung Adeno.
Lung Squamous
Melanoma
MM
Neuroblastoma
Ovarian
PDAC
Prostate
SCLC
Wilms Tumor
PCNP
CFD
SPN
CLEC12A
TRPS1
ESR1
ANKRD30A
PRLR
CRYAB
NDUFA4L2
ANGPTL4
SLC3A1
TESC
CDA
CXCL6
NPSR1
H3-5
MPL
CCDC27
AREG
CXCL3
DPEP1
CDX1
CD79A
CD79B
NCF1
TCL1A
PLTP
ELN
C9orf24
VWA3A
PTPRZ1
MT3
PMP2
OLIG1
APOA2
ALB
RBP4
APOC3
KRT14
KRT6B
CLCA2
LGALS7
S100A9
CSTA
CALML5
FDCSP
NAPSA
SFTPB
SCGB3A2
SCGB1A1
AKR1C2
ADH7
PLCXD2
CHST7
SDCBP
MLANA
PMEL
TYR
JCHAIN
TENT5C
SLAMF7
BMP6
NPY
STMN2
PHOX2A
HAND2
RHEX
CMTM7
MUC16
SOX17
MMP7
SYT8
CLDN18
KLK3
KLK2
PDLIM5
KLK4
HES6
ASCL1
INSM1
GRP
CCN2
PODXL
NPHS2
TCF21
Sensitivity
Specificity
0
0.5
1
0.2
0.6
1

**Figure S1. Cancer-type-dependent gene expression patterns. a.** Scatter plots per cancer type showing sensitivity (y axis) and specificity (x axis) of genes (points) to malignant cells, relative to other cell types, within each cancer type. Selected genes with unusually high sensitivity or specificity are labelled. **b.** Scatter plots per cancer type showing sensitivity (y axis) and specificity (x axis) of genes (points) to malignant cells in each cancer type, relative to malignant cells in other cancer types. Selected genes with unusually high sensitivity or specificity are labelled.

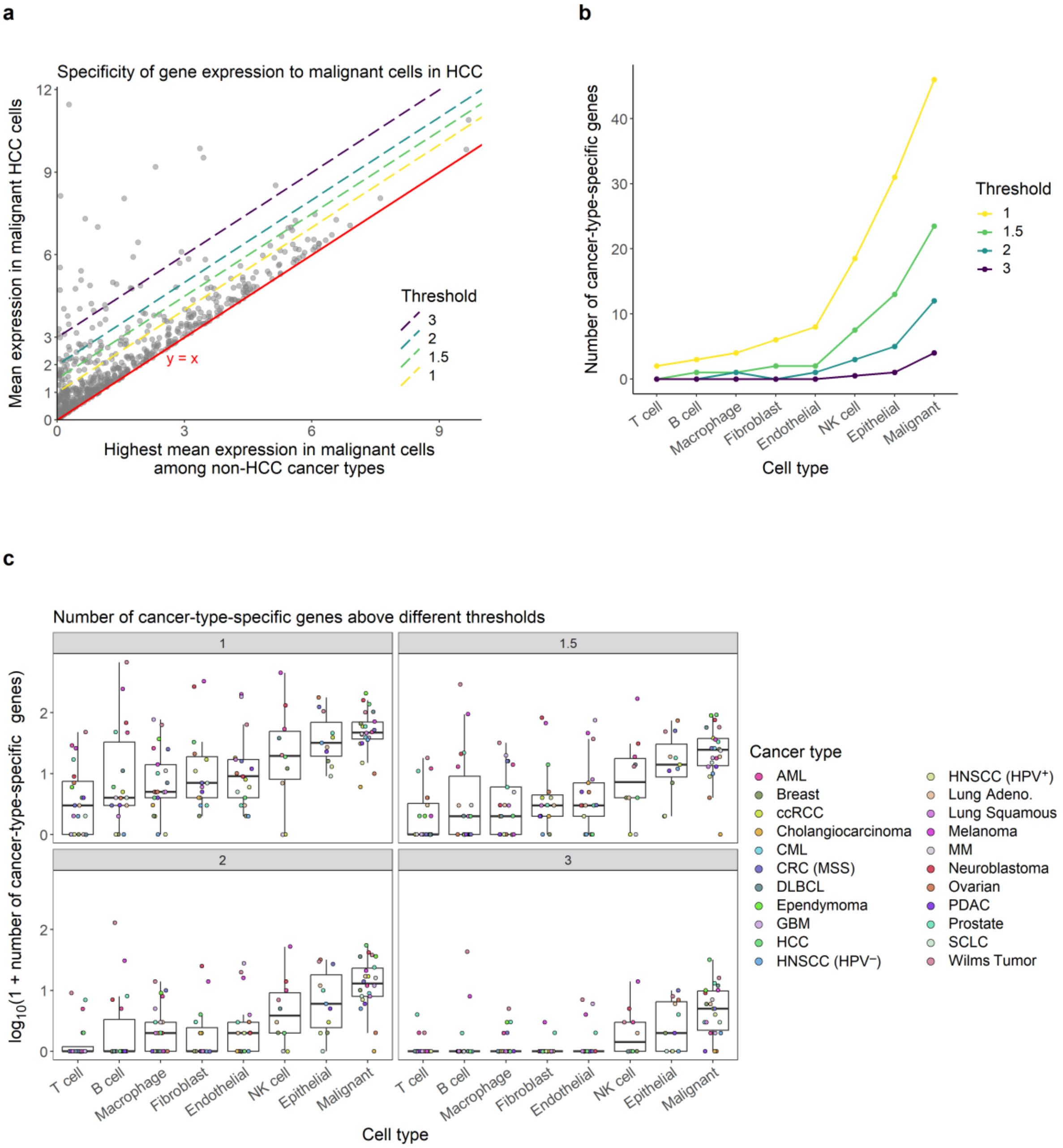


**Figure S2. Number of cancer-type-specific genes per cell type. a.** Scatter plot to illustrate the definition of cancer-type-specific gene expression at different thresholds. Points correspond to genes whose mean expression in malignant cells is highest in HCC than in all other cancer types. A point's y axis value denotes the average expression of this gene in malignant cells in HCC, while its x axis value corresponds to the maximum of its mean expression levels in malignant cells across all non-HCC cancer types. Each dashed line denotes a choice of threshold, whereby the number of genes whose expression in malignant cells is specific to HCC is defined as the number of points above this dashed line. **b.** Line plot showing the median number of cancer-type-specific genes (y axis, median across cancer types) for each cell type (x axis) for different choices of threshold (colour). Cell types are ordered by their average y values. **c.** Boxplots showing the log-transformed number of cancer-type-specific genes (y axis) per cancer type (points/colour) for each cell type (x axis), separately for each choice of threshold (panels). Cell types are ordered as in **b**. Each point in **b** corresponds to the median of points for the corresponding box in **c**, after reversing the log transformation.

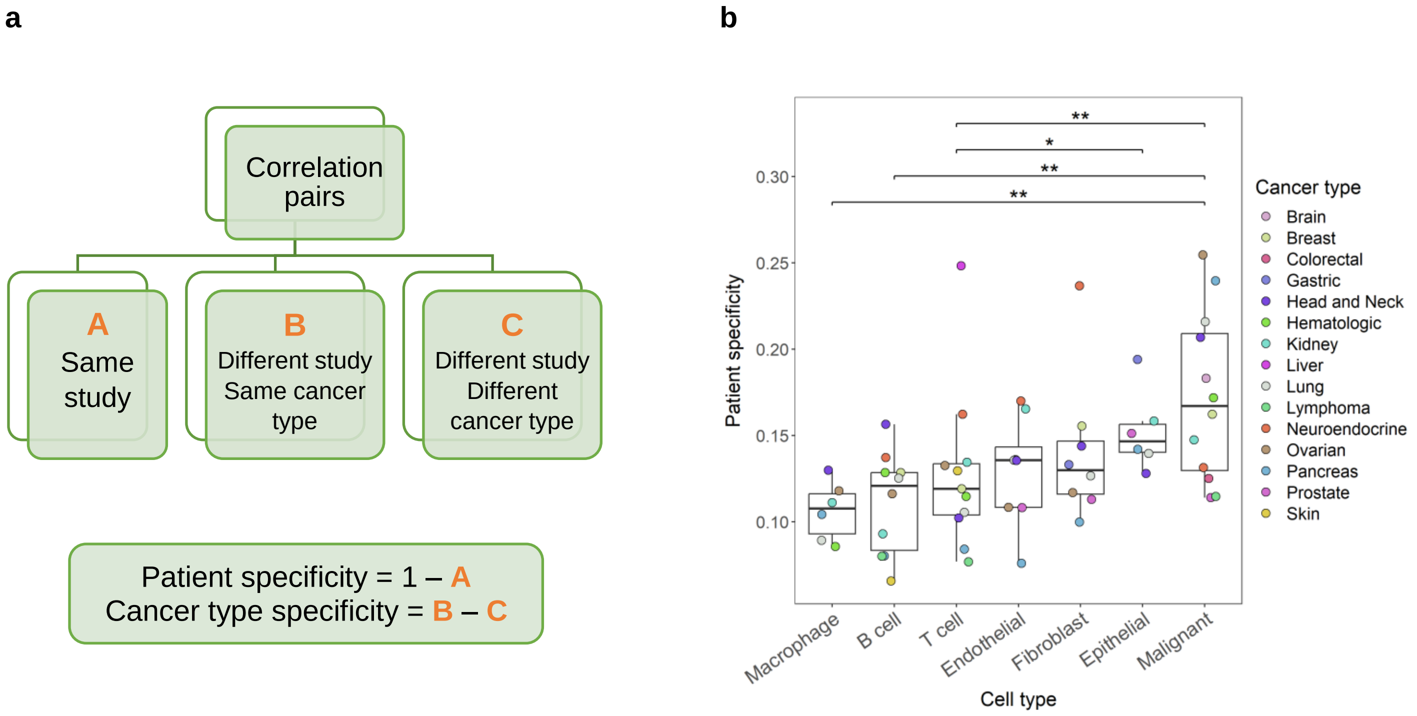


**Figure S3. Cancer type and patient specificity of cell type expression profiles. a.** Scheme illustrating the definition of cancer type and patient specificity in terms of pairwise correlations of pseudobulk profiles. **b.** Box plot showing the patient specificity (y axis) of each cell type (x axis) in each cancer type (points/colour). Asterisks indicate statistical significance (without adjustment) according to pairwise paired t tests, with '**' and '*' indicating $p < 0.01$ and $p < 0.05$ respectively. Unmarked pairwise differences are not significant.

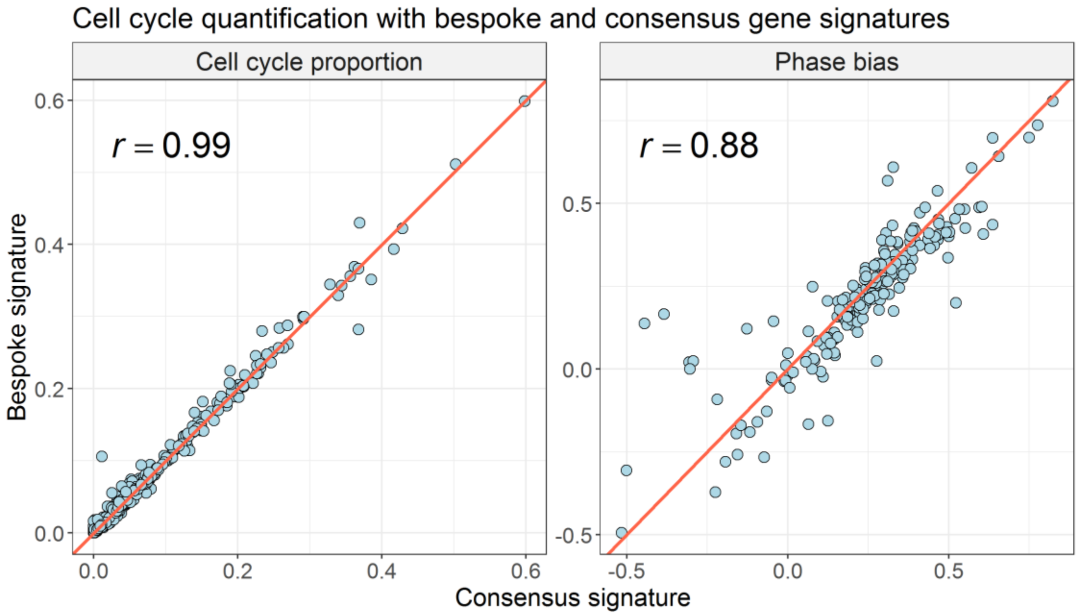


**Figure S4. Cell cycle quantification using bespoke and consensus G1/S and G2/M gene signatures.** Scatter plots showing measurements of cell cycle proportion and phase bias, respectively, in each cell type and dataset using bespoke (y axis) and consensus (x axis) G1/S and G2/M gene signatures. Each point corresponds to one cell type in one dataset. The red lines correspond to y = x, and *r* denotes Pearson correlation.

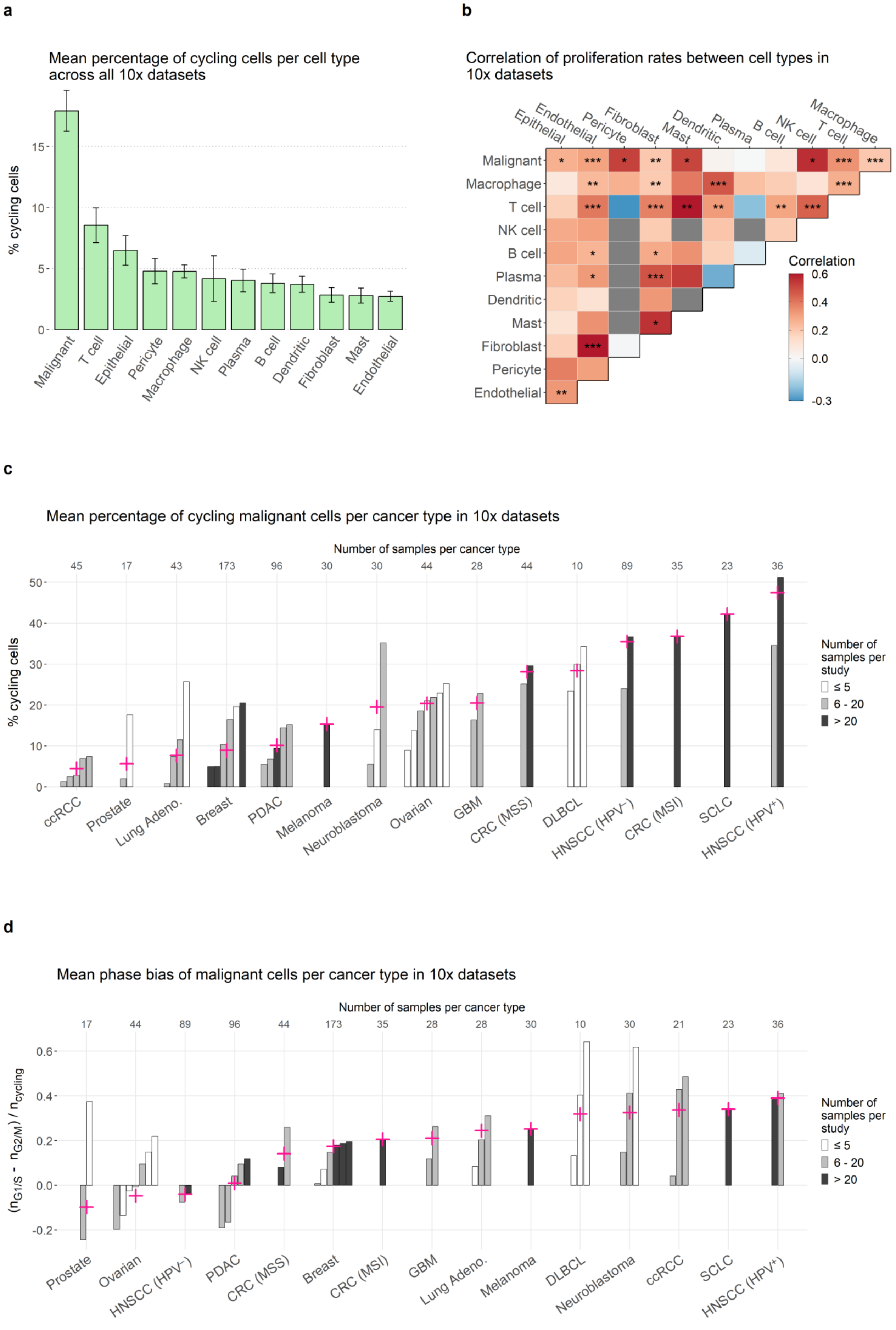

a
Mean percentage of cycling cells per cell type across all 10x datasets
% cycling cells
0
5
10
15
Malignant
T cell
Epithelial
Pericyte
Macrophage
NK cell
Plasma
B cell
Dendritic
Fibroblast
Mast
Endothelial
b
Correlation of proliferation rates between cell types in 10x datasets
Epithelial
Endothelial
Pericyte
Fibroblast
Mast
Dendritic
Plasma
B cell
NK cell
T cell
Macrophage
Malignant
Macrophage
T cell
NK cell
B cell
Plasma
Dendritic
Mast
Fibroblast
Pericyte
Endothelial
Correlation
0.6
0.4
0.2
0.0
-0.3
c
Mean percentage of cycling malignant cells per cancer type in 10x datasets
Number of samples per cancer type
45 17 43 173 96 30 30 44 28 44 10 89 35 23 36
% cycling cells
0
10
20
30
40
50
ccRCC
Prostate
Lung Adeno.
Breast
PDAC
Melanoma
Neuroblastoma
Ovarian
GBM
CRC (MSS)
DLBCL
HNSCC (HPV⁻)
CRC (MSI)
SCLC
HNSCC (HPV⁺)
Number of samples per study
≤ 5
6 - 20
> 20
d
Mean phase bias of malignant cells per cancer type in 10x datasets
Number of samples per cancer type
17 44 89 96 44 173 35 28 28 30 10 30 21 23 36
$(n_{G1/S} - n_{G2/M}) / n_{cycling}$
-0.2
0.0
0.2
0.4
0.6
Prostate
Ovarian
HNSCC (HPV⁻)
PDAC
CRC (MSS)
Breast
CRC (MSI)
GBM
Lung Adeno.
Melanoma
DLBCL
Neuroblastoma
ccRCC
SCLC
HNSCC (HPV⁺)
Number of samples per study
≤ 5
6 - 20
> 20

**Figure S5. Comparison of cell cycle patterns using only 10x data. a.** Bar plot showing the average percentage of cycling cells (y axis, average across all 10x datasets) in each cell type (x axis). Error bars denote the standard error. **b.** Heatmap showing the Spearman correlation between cell types (colour, correlation computed across samples profiled by 10x) of percentages of cycling cells. Asterisks indicate the statistical significance of the correlation (computed via algorithm AS 89[43]), after adjusting to FDR < 0.05, with '***', '**' and '*' indicating $p < 0.001$, $p < 0.01$ and $p < 0.05$ respectively, and no asterisk indicating not significant. **c.** Bar plot showing the percentage of cycling malignant cells (y axis) in each 10x dataset (bars), grouped by cancer type (x axis), with crosses denoting the average y value for each cancer type, weighted by the number of samples in each dataset which contain at least 10 malignant cells. Bar colour categorises studies by number of such samples, and values above the plot denote the total number of such samples per cancer type. **d.** Bar plot showing the phase bias (y axis, quantified by the relative fraction of cycling cells in G1/S versus G2/M) of malignant cells in each 10x dataset (bars), grouped by cancer type (x axis). Crosses, bar colour and number of samples per cancer type are defined as in **c**. Low and high y values indicate bias toward G2/M and G1/S, respectively.

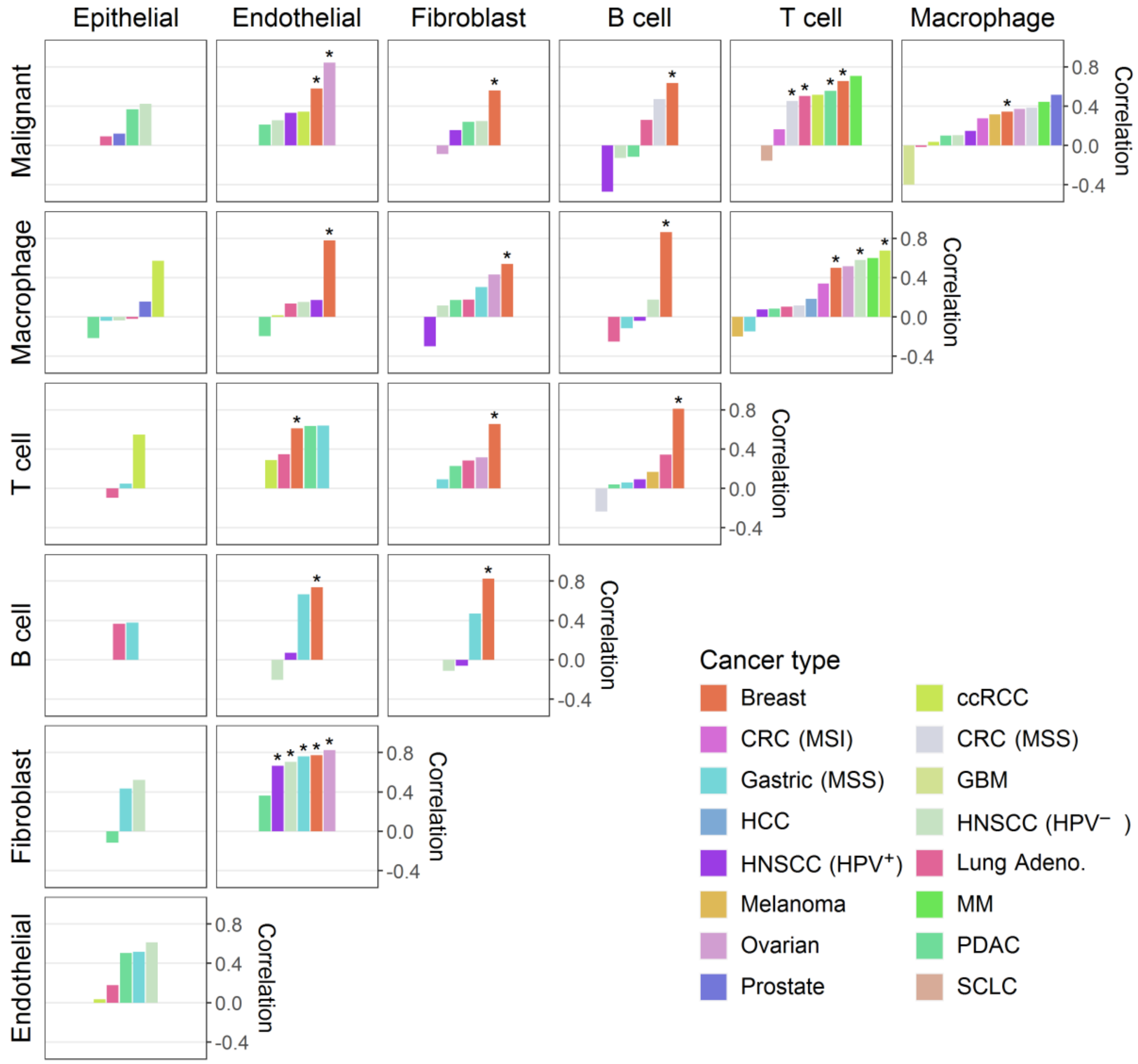


**Figure S6. Correlation of cell cycle between cell types, per cancer type.** Bar plots, for each pair of cell types, showing the Spearman correlation (across samples) of proportion of cycling cells between those cell types in each cancer type. Asterisks indicate statistical significance (adjusted p value < 0.05; p values computed via algorithm AS 89[43]; adjustment to FDR < 0.05).

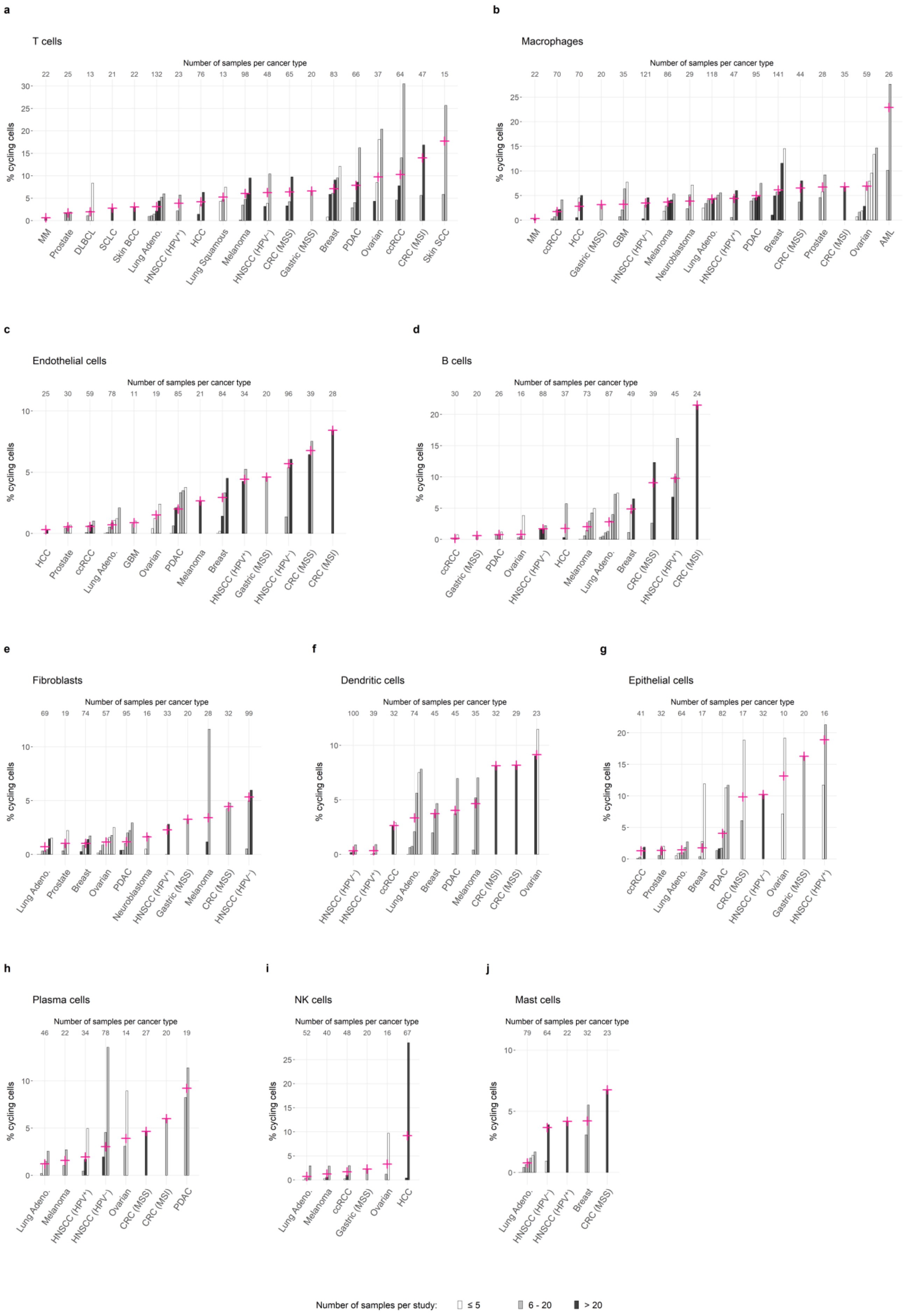

a
T cells
Number of samples per cancer type
22 25 13 21 22 132 23 76 13 98 48 65 20 83 66 37 64 47 15
% cycling cells
0 5 10 15 20 25 30
MM
Prostate
DLBCL
SCLC
Skin BCC
Lung Adeno.
HNSCC (HPV+)
HCC
Lung Squamous
Melanoma
HNSCC (HPV−)
CRC (MSS)
Gastric (MSS)
Breast
PDAC
Ovarian
ccRCC
CRC (MSI)
Skin SCC
b
Macrophages
Number of samples per cancer type
22 70 70 20 35 121 86 29 118 47 95 141 44 28 35 59 26
% cycling cells
0 5 10 15 20 25
MM
ccRCC
HCC
Gastric (MSS)
GBM
HNSCC (HPV−)
Melanoma
Neuroblastoma
Lung Adeno.
HNSCC (HPV+)
PDAC
Breast
CRC (MSS)
Prostate
CRC (MSI)
Ovarian
AML
c
Endothelial cells
Number of samples per cancer type
25 30 59 78 11 19 85 21 84 34 20 96 39 28
% cycling cells
0 5 10
HCC
Prostate
ccRCC
Lung Adeno.
GBM
Ovarian
PDAC
Melanoma
Breast
HNSCC (HPV+)
Gastric (MSS)
HNSCC (HPV−)
CRC (MSS)
CRC (MSI)
d
B cells
Number of samples per cancer type
30 20 26 16 88 37 73 87 49 39 45 24
% cycling cells
0 5 10 15 20
ccRCC
Gastric (MSS)
PDAC
Ovarian
HNSCC (HPV−)
HCC
Melanoma
Lung Adeno.
Breast
CRC (MSS)
HNSCC (HPV+)
CRC (MSI)
e
Fibroblasts
Number of samples per cancer type
69 19 74 57 95 16 33 20 28 32 99
% cycling cells
0 5 10
Lung Adeno.
Prostate
Breast
Ovarian
PDAC
Neuroblastoma
HNSCC (HPV+)
Gastric (MSS)
Melanoma
CRC (MSS)
HNSCC (HPV−)
f
Dendritic cells
Number of samples per cancer type
100 39 32 74 45 45 35 32 29 23
% cycling cells
0 5 10
HNSCC (HPV−)
HNSCC (HPV+)
ccRCC
Lung Adeno.
Breast
PDAC
Melanoma
CRC (MSI)
CRC (MSS)
Ovarian
g
Epithelial cells
Number of samples per cancer type
41 32 64 17 82 17 32 10 20 16
% cycling cells
0 5 10 15 20
ccRCC
Prostate
Lung Adeno.
Breast
PDAC
CRC (MSS)
HNSCC (HPV−)
Ovarian
Gastric (MSS)
HNSCC (HPV+)
h
Plasma cells
Number of samples per cancer type
46 22 34 78 14 27 20 19
% cycling cells
0 5 10
Lung Adeno.
Melanoma
HNSCC (HPV+)
HNSCC (HPV−)
Ovarian
CRC (MSS)
CRC (MSI)
PDAC
i
NK cells
Number of samples per cancer type
52 40 48 20 16 67
% cycling cells
0 5 10 15 20 25
Lung Adeno.
Melanoma
ccRCC
Gastric (MSS)
Ovarian
HCC
j
Mast cells
Number of samples per cancer type
79 64 22 32 23
% cycling cells
0 5 10
Lung Adeno.
HNSCC (HPV−)
HNSCC (HPV+)
Breast
CRC (MSS)
Number of samples per study:
≤ 5
6 - 20
> 20

**Figure S7. Proliferation rates across cancer types for non-malignant cell types. a-k.** Bar plots, for each of the most common non-malignant cell types, showing the percentage of cycling cells of that type (y axis) in each study (bars), grouped by cancer type (x axis), with crosses denoting the average y value for each cancer type, weighted by the number of samples in each study which contain at least 10 cells of that type. Bar colour categorises studies by number of such samples, and values above the plot denote the total number of such samples in each cancer type.

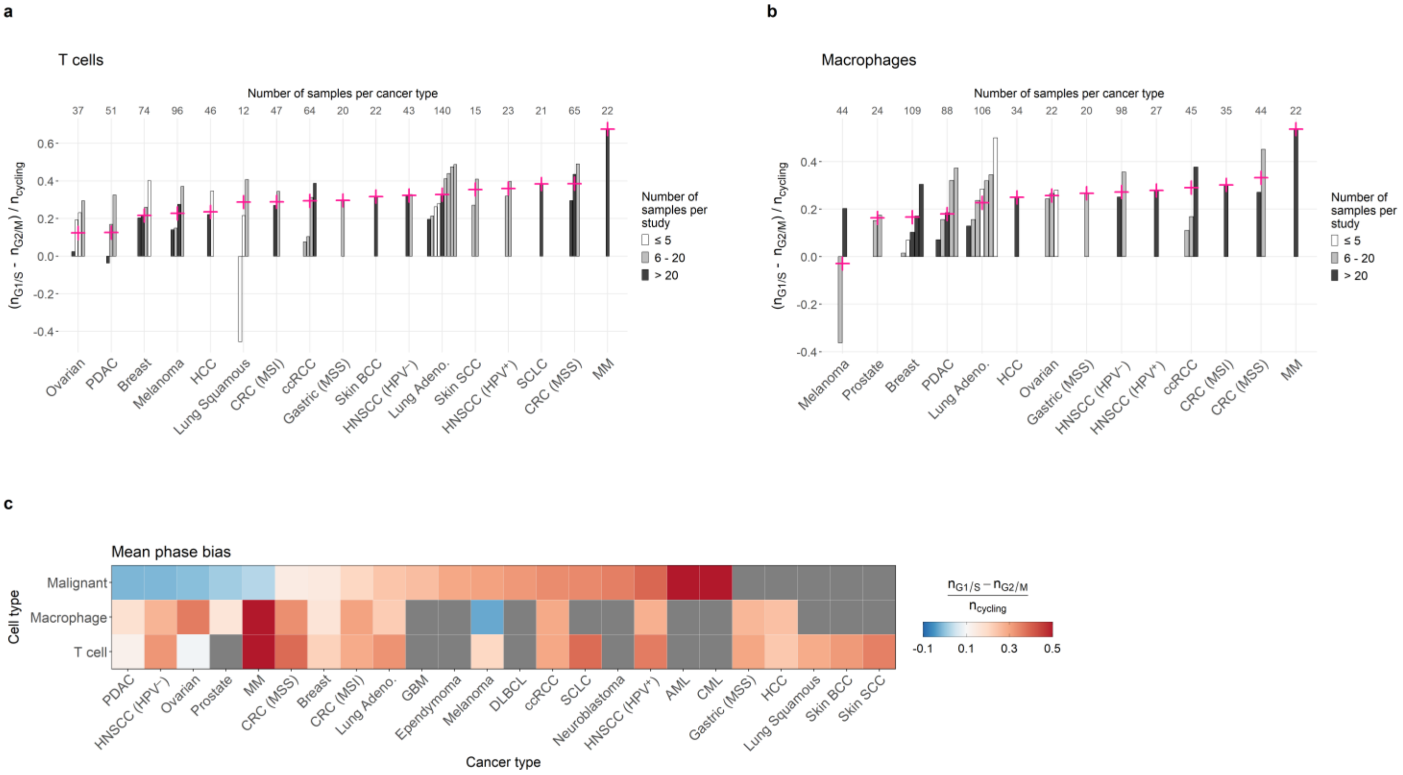

**Figure S8. Phase bias patterns across cancer types. a.** Bar plot showing the phase bias (y axis, quantified by the relative fraction of cycling cells in G1/S versus G2/M) of T cells in each study (bars), grouped by cancer type (x axis), with crosses denoting the average y value for each cancer type, weighted by the number of samples in each study which contain at least 10 T cells. Bar colour categorises studies by number of such samples, and values above the plot denote the total number of such samples in each cancer type. Low and high y values indicate bias toward G2/M and G1/S, respectively. **b.** Bar plot as in **a** for macrophages. **c.** Heatmap showing the weighted average of the phase bias (colour, defined as for the crosses in **a**) per cancer type (x axis) and cell type (y axis). Grey squares indicate insufficient data.

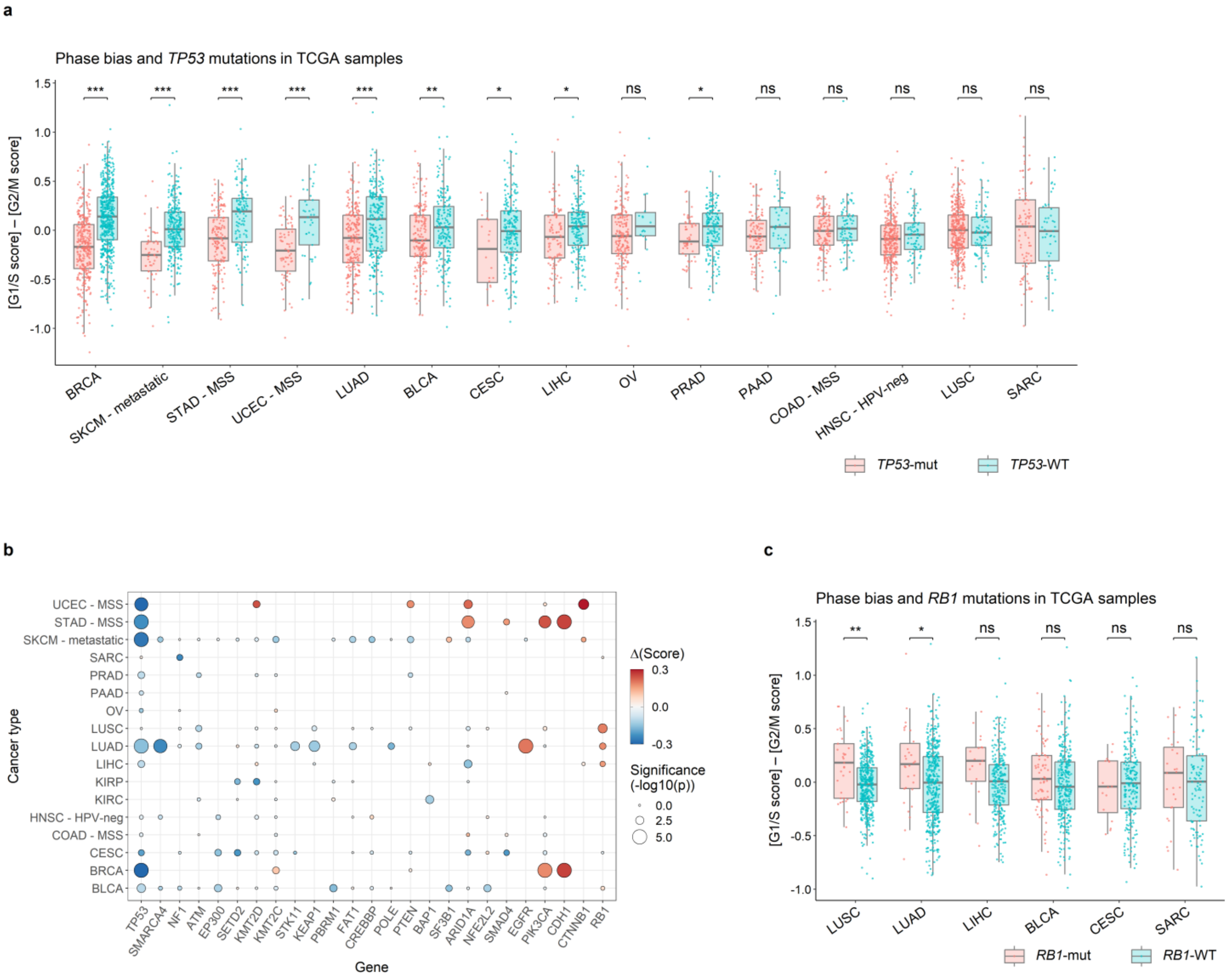


**Figure S9. Genomic associations of phase bias. a.** Box plot showing the phase bias scores (y axis, defined as the difference between scores for G1/S and G2/M gene signatures) of TCGA tumour samples (points), grouped by cancer type (x axis) and coloured by *TP53* mutation status. Significance values were computed by t test and adjusted to FDR < 0.05. Asterisks indicate significance after adjustment in each case, with '***', '**' and '*' indicating p < 0.001, p < 0.01 and p < 0.05 respectively. Low and high y values indicate bias toward G2/M and G1/S, respectively. **b.** Dot plot showing, for a selection of genes commonly mutated in cancer (x axis), the difference in average phase bias score between mutant and wild-type tumours (point colour; phase bias score defined as in **a**) and the statistical significance of this difference (point size, computed as in **a**, before adjustment) in each cancer type (y axis). **c.** Box plot as in **a** for *RB1* mutations.
